## Supplementary data for "Targeted Protein Degradation in *Escherichia coli* - depletion of the essential GroEL using CLIPPERs"

### Supplementary files

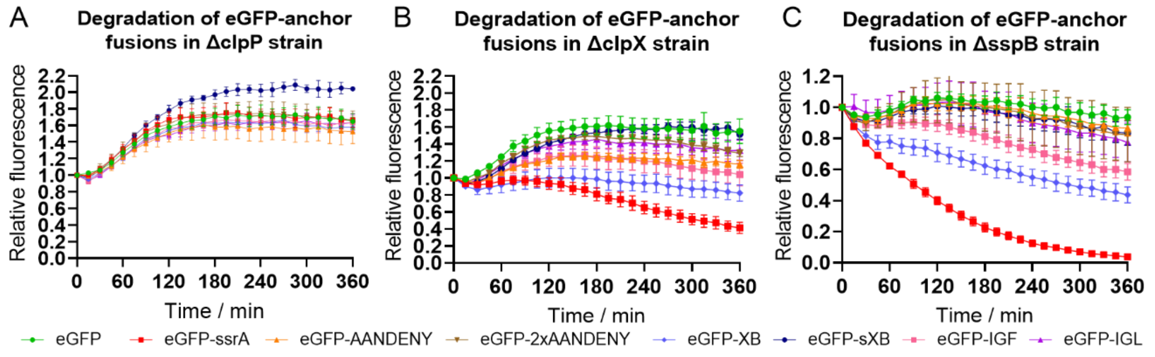


**Supplementary Figure S1. Degradation of the eGFP-anchor fusion proteins in bacteria.** Degradation was performed in *E. coli* deletion mutant strains (A) ΔclpP, (B) ΔclpP, and (C) ΔsspB.

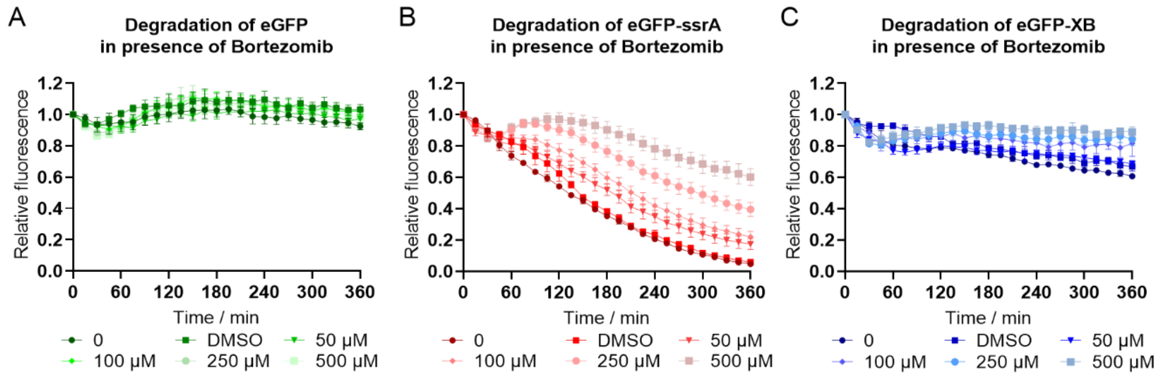


**Supplementary Figure S2. Degradation of the eGFP-anchor fusion proteins in bacteria in presence of bortezomib.** Degradation was performed in *E. coli* BW25113 strain for (A) untagged eGFP (negative control), (B) eGFP-ssrA (positive control), and (C) eGFP-XB.

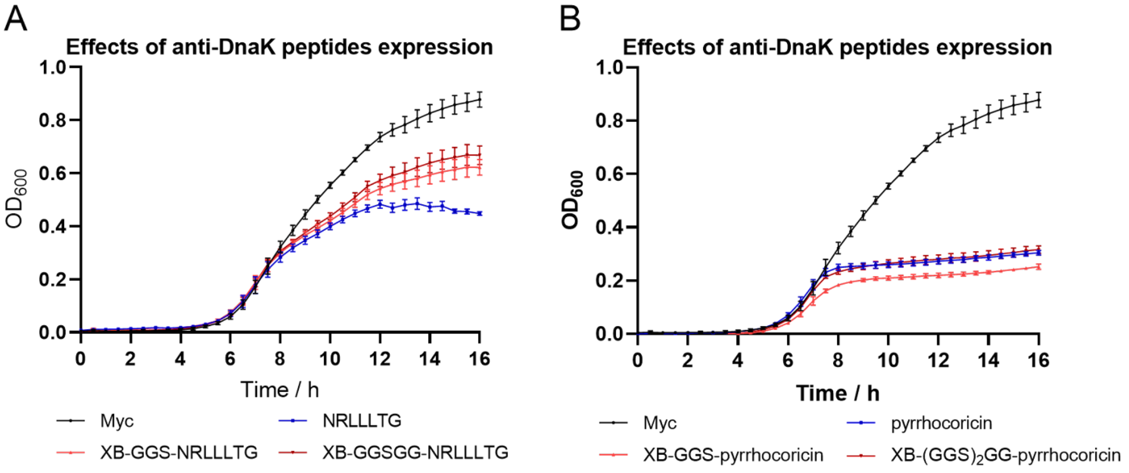


**Supplementary Figure S3. Growth curves of bacteria grown at 30 °C in presence of expression-inducing arabinose.** The assay was performed for bacteria transformed with pBAD-Myc plasmids encoding DnaK-targeting CLIPPERs with (A) NRLLLTG peptide as bait, or (B) pyrrhocoricin peptide as bait.

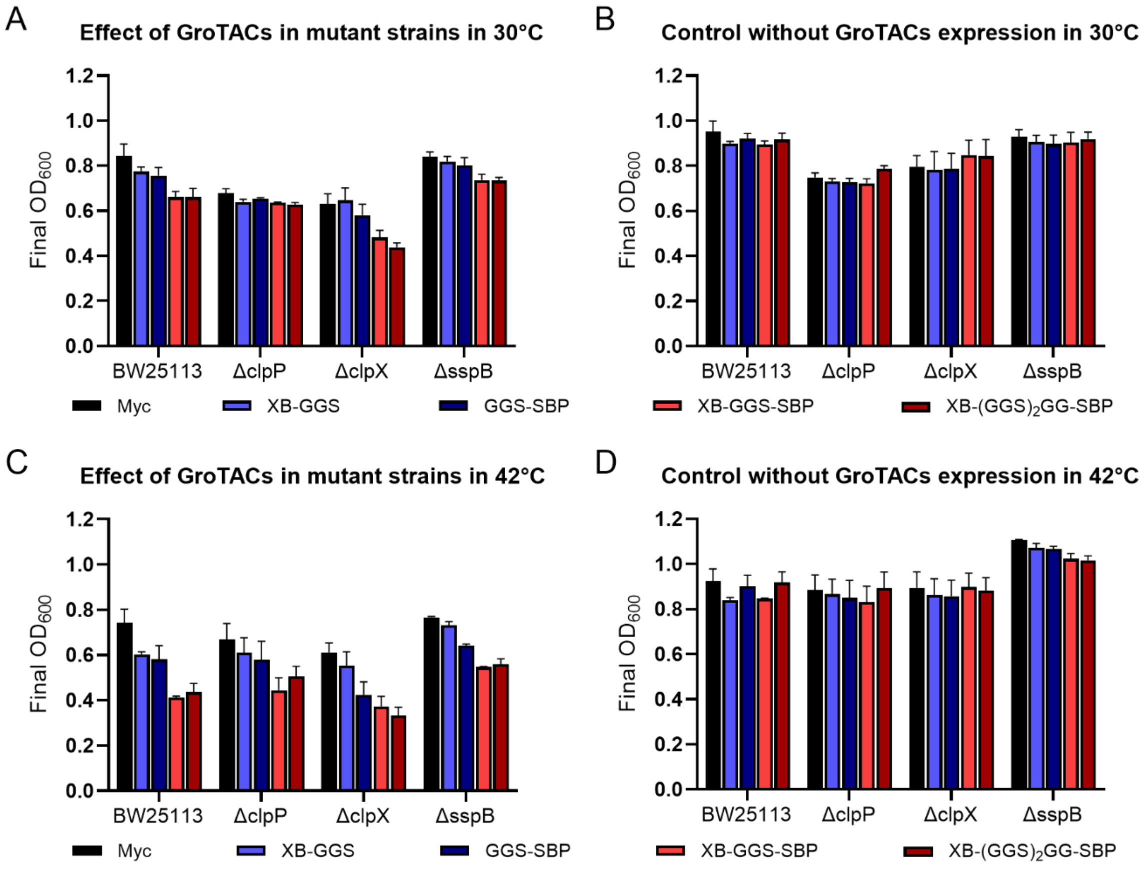


**Supplementary Figure S4. The effect of protease component mutations on the effect of GroTACs on growth inhibition.** The expression of GroTACs was tested in *E. coli* deletion mutant strains for their effect on final culture $OD_{600}$ after 16 h of culturing (A and C) in presence, or (B and D) in absence of expression-inducing arabinose at 30 °C (A and B) or 42 °C (C and D).

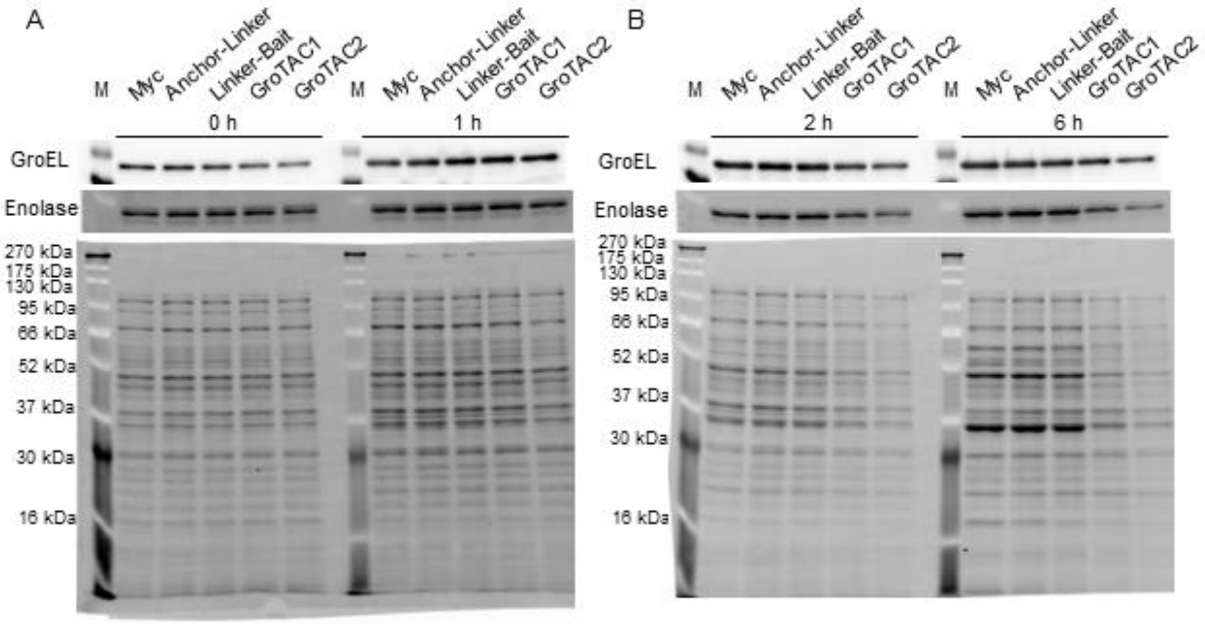


**Supplementary Figure S5. The effect or GroTACs expression on proteins in *E. coli*.** The figure presents representative western blots of GroEL and enolase levels and total protein on PVDF membranes visualized by stain-free method [68]. Protein levels were measured (A) 0 h and 1 h after induction and (B) 2 h and 6 h after induction. (M – protein size marker)

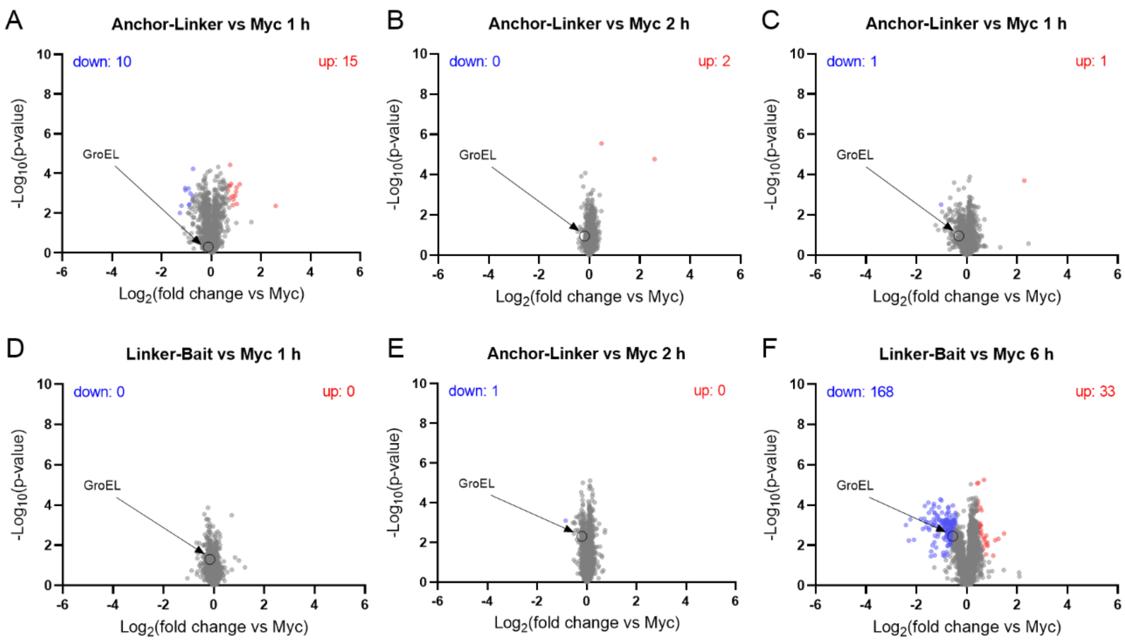


**Supplementary Figure S6. Changes of protein levels upon expression of control peptides.** (A) – (C) Volcano plots picturing the effects of Anchor-Linker peptide in different time points. (D) – (F) Volcano plots picturing the effects of Linker-Bait peptide in different time points. GroEL is indicated with an arrow and a circle. Significantly down-regulated proteins are marked in blue and up-regulated proteins in red.

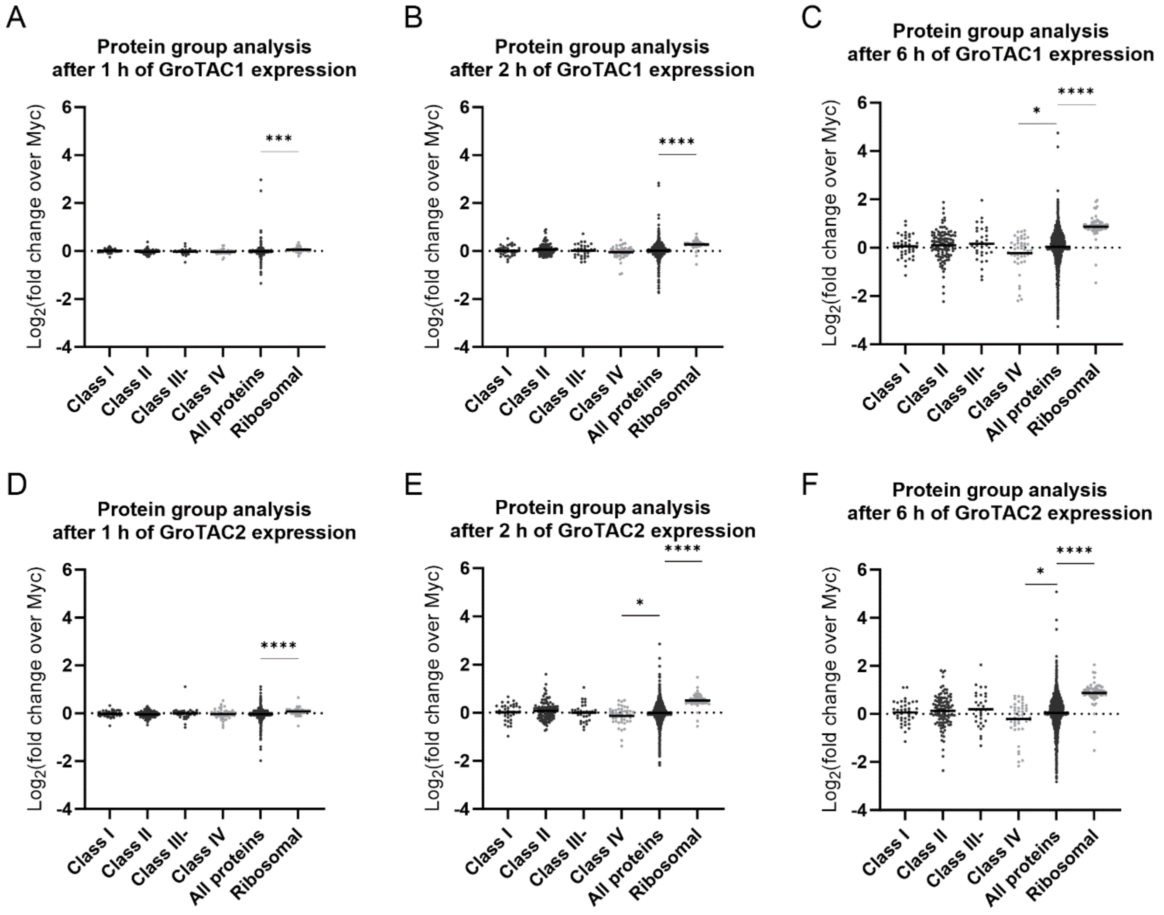


**Supplementary Figure S7. GroTAC-induced changes in specific protein groups** (Group analysis of relative changes in levels of GroEL substrates and ribosomal proteins induced by GroTAC1 (A-C) and GroTAC2 (D-F) after 1 h (A and D) 2 h (B and E) and 6 h (C and F) of peptide expression. Statistically significant changes are marker with asterisks (* p-val < 0.05; *** p-val < 0.001; **** p-val < 0.0001).

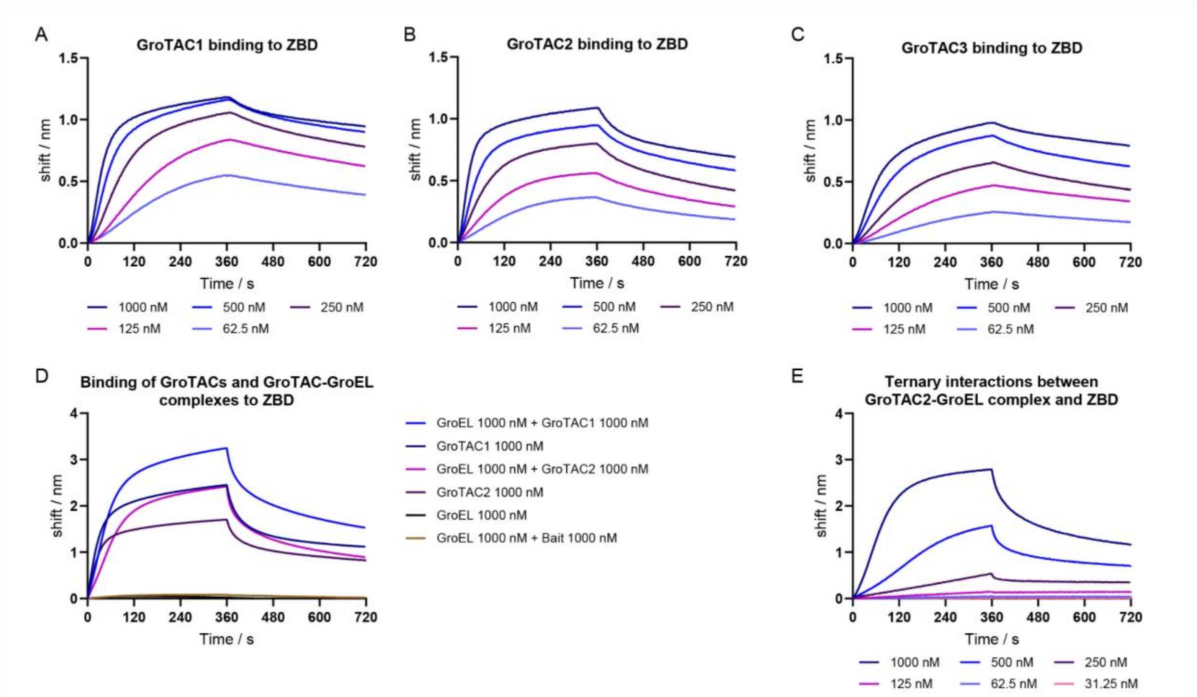


**Supplementary Figure S8. Binding of untagged GroTACs to ZBD measured by BLI.** (A) Binding of GroTAC1 to immobilised His-ZBD; (B) Binding of GroTAC2 to immobilised His-ZBD; (C) Binding of GroTAC3 to immobilised His-ZBD; (D) Comparison of binding of GroEL, GroTACs or their complexes to immobilized ZBD. Bait is a control peptide consisting of a linker-SBP fusion. (E) Binding of GroEL-GroTAC2 complex to immobilised His-ZBD using fixed GroEL concentration (1000 nM) and increasing GroTAC2 concentrations.

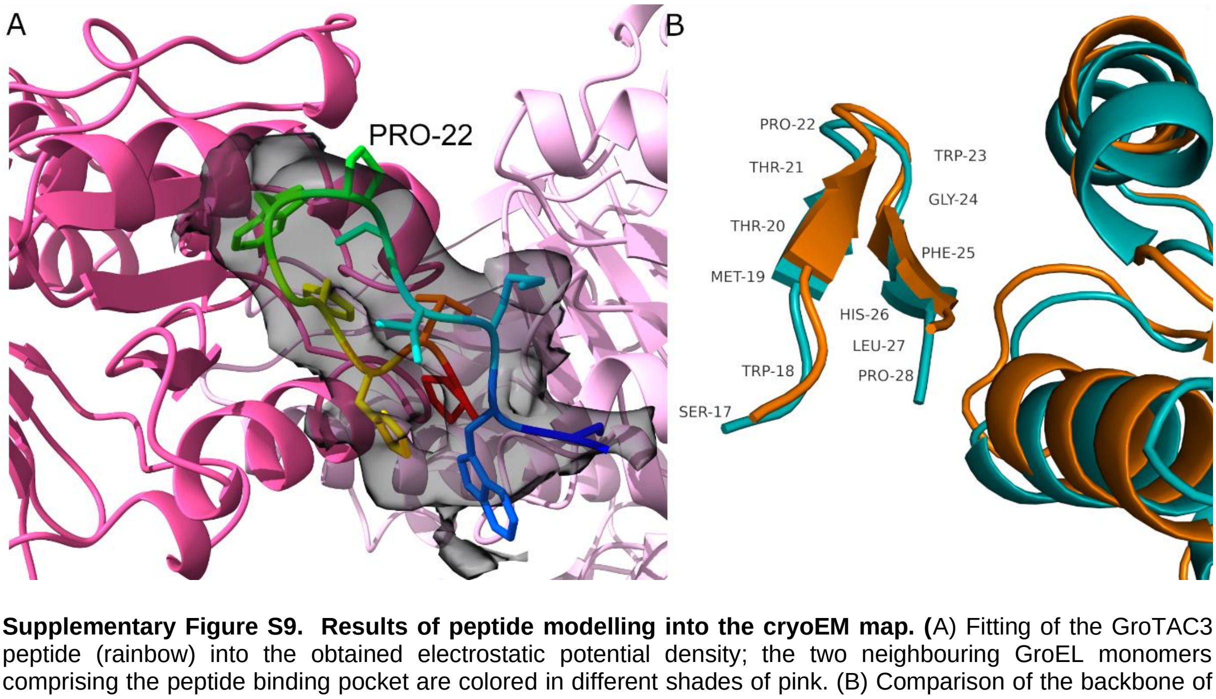


**Supplementary Figure S9. Results of peptide modelling into the cryoEM map. (**A) Fitting of the GroTAC3 peptide (rainbow) into the obtained electrostatic potential density; the two neighbouring GroEL monomers comprising the peptide binding pocket are colored in different shades of pink. (B) Comparison of the backbone of GroEL-binding part of GroTAC peptide obtained from CryoEM (cyan; PDB ID 8S32) with the crystallographic structure of SBP in complex with GroEL (orange, PDB ID 1MNF).

**Table S1. Binding parameters between peptides and their binding partners ClpX and GroEL.** The results were calculated from BLI experiments presented in Figure 4 and Supplementary Figure S8. The parameters were calculated assuming 1:1 binding stoichiometry to avoid overfitting of the data.

| Analyte | Ligand | $K_D$ (M) | $k_a$ (1/Ms) | $k_{dis}$ (1/s) | Full $X^2$ | Full $R^2$ |
|---|---|---|---|---|---|---|
| ClpX | His-SUMO-XB-GGS | 6.48E-08 ± 2.69E-10 | 7.65E+04 ± 2.83E+02 | 4.95E-03 ± 9.39E-06 | 12.878 | 0.996 |
| ClpX | His-SUMO-XB-GGS-SBP | 7.04E-08 ± 5.32E-10 | 1.09E+05 ± 7.72E+02 | 7.69E-03 ± 2.06E-05 | 13.202 | 0.987 |
| GroEL | His-SUMO-GGS-SBP | 9.36E-08 ± 1.37E-09 | 9.82E+04 ± 1.36E+03 | 9.19E-03 ± 4.11E-05 | 47.844 | 0.931 |
| GroEL | His-SUMO-XB-GGS-SBP | 1.54E-07 ± 1.45E-09 | 2.66E+04 ± 2.21E+02 | 4.09E-03 ± 1.84E-05 | 8.5745 | 0.959 |
| GroTAC1 | His-ZBD | 3.54E-08 ± 1.67E-10 | 2.26E+04 ± 6.70E+01 | 7.98E-04 ± 2.92E-06 | 19.622 | 0.989 |
| GroTAC2 | His-ZBD | 5.26E-08 ± 2.84E-10 | 3.14E+04 ± 1.40E+02 | 1.65E-03 ± 5.02E-06 | 27.907 | 0.980 |
| GroTAC3 | His-ZBD | 8.24E-08 ± 4.59E-10 | 1.11E+04 ± 4.30E+01 | 9.15E-04 ± 3.65E-06 | 14.097 | 0.989 |
| GroEL-GroTAC1 complex | His-ZBD | 1.65E-07 ± 1.71E-09 | 1.09E+04 ± 9.32+01 | 1.80E-03 ± 1.05E-05 | 295.552 | 0.973 |
| GroEL-GroTAC2 complex | His-ZBD | 3.08E-07 ± 3.79E-09 | 9.32E+03 ± 1.05E+02 | 2.87E-03 ± 1.45E-05 | 238.550 | 0.963 |

**Table S2. List of DNA constructs used in the study**

| Construct name | Purpose | Backbone | Insert | Source |
|---|---|---|---|---|
| pBAD-6xHis-SUMO-ClpX | expression and purification of ClpX | pBAD | His tag - SUMO tag - ClpX | [33] |
| pET28a-6xHis-SUMO-ClpP | expression and purification of ClpP | pET28a | His tag - SUMO tag - ClpP | [33] |
| pET28a-6xHis-SUMO-SspB | expression and purification of SspB | pET28a | His tag - SUMO tag - SspB | [33] |
| pET28a-His-ZBD | expression and purification of ZBD | pET28a | His tag - ZBD | this study |
| pET28a-GroEL | expression and purification of GroEL | pET28a | GroEL | this study |
| pET28a-6xHis-SUMO-XB-GGS | expression and purification of anchor-linker | pET28a | His tag - SUMO tag - CYRGGRPALRVVK - GGS | this study |

| | | | | |
|---|---|---|---|---|
| pET28a-6xHis-SUMO-GGS-SBP | expression and purification of linker bait against GroEL | pET28a | His tag - SUMO tag - CYRGGRPALRVVKSBP | this study |
| pET28a-6xHis-SUMO-XB-GGS-SBP | expression and purification of GroTAC1 | pET28a | His tag - SUMO - GGS - SWMTTPWGFHLP | this study |
| pBAD-His-TEV-cAbGFP | expression and purification of | pBAD | His tag - TEV cleavage site - cAbGFP | this study |
| pBAD-6xHIS-TEV | eGFP-anchor stability assay | pBAD | His tag - T7 gene leader - TEV cleavage site | [33] |
| pBAD-6xHis-TEV-eGFP | eGFP-anchor stability assay | pBAD | His tag - T7 gene leader - TEV cleavage site - eGFP | Addgene #54762 |
| pBAD-6xHis-TEV-eGFP-ssrA | eGFP-anchor stability assay | pBAD | His tag - T7 gene leader - TEV cleavage site - eGFP - AANDENYALAA | [33] |
| pBAD-6xHis-TEV-eGFP-AANDENY | eGFP-anchor stability assay | pBAD | His tag - T7 gene leader - TEV cleavage site - eGFP - AANDENY | [33] |
| pBAD-6xHis-TEV-eGFP-2xAANDENY | eGFP-anchor stability assay | pBAD | His tag - T7 gene leader - TEV cleavage site - eGFP - AANDENYAANDENY | [33] |
| pBAD-6xHis-TEV-eGFP-XB | eGFP-anchor stability assay | pBAD | His tag - T7 gene leader - TEV cleavage site - eGFP - CYRGGRPALRVVK | this study |
| pBAD-6xHis-TEV-eGFP-sXB | eGFP-anchor stability assay | pBAD | His tag - T7 gene leader - TEV cleavage site - eGFP - ALRVVK | this study |
| pBAD-6xHis-TEV-eGFP-IGF | eGFP-anchor stability assay | pBAD | His tag - T7 gene leader - TEV cleavage site - eGFP - GIGFGATVK | this study |
| pBAD-6xHis-TEV-eGFP-IGL | eGFP-anchor stability assay | pBAD | His tag - T7 gene leader - TEV cleavage site - eGFP - KSIGLIHQD | this study |
| pBAD-myc | testing CLIPPERs in bacteria (control) | pBAD | Myc tag | this study |
| pBAD-myc-XB-GGS | testing CLIPPERs in bacteria (anchor control) | pBAD | Myc tag - CYRGGRPALRVVK - GGS | this study |
| pBAD-myc-GGS-SBP | testing CLIPPERs in bacteria (GroEL bait control) | pBAD | Myc tag - GGS - SWMTTPWGFHLP | this study |
| pBAD-myc-XB-GGS-SBP | testing CLIPPERs in bacteria (GroTAC1) | pBAD | Myc tag - CYRGGRPALRVVK - GGS - SWMTTPWGFHLP | this study |
| pBAD-myc-XB-GGSGGSGG-SBP | testing CLIPPERs in bacteria (GroTAC2) | pBAD | Myc tag - CYRGGRPALRVVK - GGSGGSGG - SWMTTPWGFHLP | this study |
| pBAD-myc-NRLLLTG | testing CLIPPERs in bacteria (DnaK bait control) | pBAD | Myc tag - NRLLLTG | this study |

| | | | | |
|---|---|---|---|---|
| pBAD-myc-XB-GGS-NRLLLTG | testing CLIPPERs in bacteria (DnaK degrader 1) | pBAD | Myc tag - CYRGGRPALRVVK - GGS - NRLLLTG | this study |
| pBAD-myc-XB-GGSGG-NRLLLTG | testing CLIPPERs in bacteria (DnaK degrader 2) | pBAD | Myc tag - CYRGGRPALRVVK - GGSGG - NRLLLTG | this study |
| pBAD-myc-pyrrhocoricin | testing CLIPPERs in bacteria (pyrrhocoricin control) | pBAD | Myc tag - VDKGSYLPRPTPPRPIYNRN | this study |
| pBAD-myc-XB-GGS-pyrrhocoricin | testing CLIPPERs in bacteria (pyrrhocoricin degrader 1) | pBAD | Myc tag - CYRGGRPALRVVK - GGS - VDKGSYLPRPTPPRPIYNRN | this study |
| pBAD-myc-XB-GGSGGSGG-pyrrhocoricin | testing CLIPPERs in bacteria (pyrrhocoricin degrader 2) | pBAD | Myc tag - CYRGGRPALRVVK - GGSGGSGG - VDKGSYLPRPTPPRPIYNRN | this study |

**Table S3. List of oligonucleotides used for obtaining DNA constructs**

| **Plasmid name** | **Template** | **Forward primer** | **Reverse primer** |
|---|---|---|---|
| pBAD-eGFP-XB | pBAD-eGFP | GCATTACGCGTTGTGAAGTAAGAATTCGAAGCTTGGCTG | CGGTCGACCACCGCGGTAGCACTTGTACAGCTCGTCCATG |
| pBAD-eGFP-sXB | pBAD-eGFP | TGTGAAGTAAGAATTCGAAGCTTGGCTG | ACGCGTAAGGCCTTGTACAGCTCGTCCATG |
| pBAD-eGFP-IGF | pBAD-eGFP | CGCGACGGTAAAATAAGAATTCGAAGCTTGGC | CCAAAACCAATGCCCTTGTACAGCTCGTCCATG |
| pBAD-eGFP-IGL | pBAD-eGFP | TATCCACCAGGATTAAGAATTCGAAGCTTGGC | AGACCAATGGATTTCTTGTACAGCTCGTCCATG |
| pBAD-Myc-eGFP | pBAD-eGFP | AGCGAAGAAGATCTGGGCTCGAGCATGGTGAGC | AATCAGTTTCTGTTCCATATGTATATCTCCTTCTTAAAGTTAAACAAAATTATTTCTAG |
| pBAD-Myc-eGFP-XB | pBAD-eGFP-XB | AGCGAAGAAGATCTGGGCTCGAGCATGGTGAGC | AATCAGTTTCTGTTCCATATGTATATCTCCTTCTTAAAGTTAAACAAAATTATTTCTAG |
| pBAD-Myc | pBAD-Myc-eGFP | TAAGAATTCGAAGCTTGGC | GCTCGAGCCCAGATCTTC |
| pBAD-Myc-XB | pBAD-Myc-eGFP-XB | TGCTACCGCGGTGGTCGA | GCTCGAGCCCAGATCCTCTTC |
| pBAD-Myc-XB-GGS-SBP | pBAD-Myc-XB-GGS-SSB | TGGGGTTTTCACCTGCCCTAAGAATTCGAAGCTTGGC | AGGCGTAGTCATCCACGAGCTACCGCCCTTCACAAC |
| pBAD-Myc-XB-GGS | pBAD-Myc-XB-GGS-SSB | TAAGAATTCGAAGCTTGGC | GCTACCGCCCTTCACAAC |
| pBAD-Myc-GGS-SBP | pBAD-Myc-XB-GGS-SBP | GGCGGTAGCTCGTGGATG | GCTCGAGCCCAGATCCTCTTC |

| | | | |
|---|---|---|---|
| pBAD-Myc-XB-GGSGGSGG-SBP | pBAD-Myc-XB-GGSGGSGG-SSB | TGGGGTTTTCACCTGCCCTAAGAATTCGAAGCTTGGC | AGGCGTAGTCATCCACGAACCGCCGCTACCGCCGCT |
| pBAD-Myc-NRLLLTG | pBAD-Myc | GCTGACTGGTTAAGAATTCGAAGCTTGGCTG | AGCAGACGGTTGCTCGAGCCCAGATCTTC |
| pBAD-Myc-XB-GGS-NRLLLTG | pBAD-Myc-XB-GGS-SSB | GCTGACTGGTTAAGAATTCGAAGCTTGGCTG | AGCAGACGGTTGCTACCGCCCTTCACAAC |
| pBAD-Myc-XB-GGSGG-NRLLLTG | pBAD-Myc-XB-GGSGG-SSB | GCTGACTGGTTAAGAATTCGAAGCTTGGCTG | AGCAGACGGTTACCGCCGCTACCGCCGCT |
| pBAD-Myc-pyrrhocoricin | pBAD-Myc | ACGCCACCACGCCCCATCTACAACCGTAATTAAGAATTCGAAGCTTGGCTG | CGGGCGCGGTAAGTAACTGCCCTTATCCACGCTCGAGCCCAGATCTTC |
| pBAD-Myc-XB-GGS-pyrrhocoricin | pBAD-Myc-XB-GGS-SSB | ACGCCACCACGCCCCATCTACAACCGTAATTAAGAATTCGAAGCTTGGCTG | CGGGCGCGGTAAGTAACTGCCCTTATCCACGCTACCGCCCTTCACAAC |
| pBAD-Myc-XB-GGSGGSGG-pyrrhocoricin | pBAD-Myc-XB-GGSGGSGG-SSB | ACGCCACCACGCCCCATCTACAACCGTAATTAAGAATTCGAAGCTTGGCTG | CGGGCGCGGTAAGTAACTGCCCTTATCCACACCGCCGCTACCGCCGCT |
| pET28a-HisSUMO-GroEL | Genomic DNA from *E. coli* Top10 | TGATTGAAGTCTACCAGGAACAAACCGGTGGATCCATGGCAGCTAAAGACGTAAAATTCG | CGGATCTCAGTGGTGGTGGTGGTGGTGCTCGAGTTACATCATGCCGCCCATGCC |
| pET28a-GroEL | pET28a-His SUMO-GroEL | ATGGCAGCTAAAGACGTA | GGTATATCTCCTTCTTAAAGTTAAAC |
| pET28a-HisSUMO-ZBD | pET28a-SUMO-ClpX | TGATTGAAGTCTACCAGGAACAAACCGGTGGATCCACAGATAAACGCAAAGATGGCT | CGGATCTCAGTGGTGGTGGTGGTGGTGCTCGAGTTAAAATCTCTTCGCGAATGATGTCG |
| pET28a-His-ZBD | pET28a-HisSUMO-ZBD | GGTGGATCCACAGATAAACG | GTGATGATGATGATGATGG |
| pET28a-HisSUMO-XB-GGS | - (annealed oligos used as insert) | CCGGTGGATCCTGCTACCGCGGTGGTCGACCGGCATTACGCGTTGTGAAGGGCGGTAGCTAACTCGAGCACC | GGTGCTCGAGTTAGCTACCGCCCTTCACAACGCGTAATGCCGGTCGACCACCGCGGTAGCAGGATCCACCGG |
| pET28a-HisSUMO-GGS-SBP | - (annealed oligos used as insert) | CCGGTGGATCCGGCGGTAGCTCGTGGATGACTACGCCTTGGGGTTTTCACCTGCCCTAACTCGAGCACC | GGTGCTCGAGTTAGGGCAGGTGAAAACCCCAAGGCGTAGTCATCCACGAGCTACCGCCGGATCCACCGG |
| pET28a-HisSUMO-XB-GGS-SBP | - (annealed oligos used as insert) | CCGGTGGATCCTGCTACCGCGGTGGTCGACCGGCATTACGCGTTGTGAAGGGCGGTAGCTCGTG | CGTAATGCCGGTCGACCACCGCGGTAGCAGGATCCACCGG |
| | | GATGACTACGCCTTGGGGTTTTCACCTGCCCTAACTCGAGCACC | GGTGCTCGAGTTAGGGCAGGTGAAAACCCCAAGGCGTAGTCATCCACGAGCTACCGCCCTTCACAACG |

**Table S4. List of *E. coli* strains used in the study**

| Strain | Genotype | Source |
|---|---|---|

| | | |
|---|---|---|
| Top10 | F⁻ mcrA DE(mrr-hsdRMS-mcrBC) φ80lacZ DE(M15) DE(lacX)74 recA1 araD139 DE(ara-leu)7697 galU galK λ⁻ rpsL($Str^R$) endA1 nupG | Laboratory strain collection |
| BL21 (DE3) | F⁻ ompT $hsdS_B$ ($r_B^-$, $m_B^-$) gal dcm (DE3) | Laboratory strain collection |
| C43 (DE3) | F⁻ ompT $hsdS_B$ ($r_B^-$, $m_B^-$) gal dcm (DE3) $lacUV5^p$ | Laboratory strain collection |
| BW25113 | F⁻ rrnB DElacZ4787 HsdR514 DE(araBAD)567 DE(rhaBAD)568 rph-1 | Keio collection [67] |
| J42W07 | BW25113 DE(clpP)::kan | Keio collection [67] |
| JW0428 | BW25113 DE(clpX)::kan | Keio collection [67] |
| JW0866 | BW25113 DE(sspB)::kan | Keio collection [67] |
